# Towards mouse genetic-specific RNA-sequencing read mapping

**DOI:** 10.1101/2021.10.01.462776

**Authors:** Nastassia Gobet, Maxime Jan, Paul Franken, Ioannis Xenarios

## Abstract

Genetic variations affect behavior and cause disease but understanding how these variants drive complex traits is still an open question. A common approach is to link the genetic variants to intermediate molecular phenotypes such as the transcriptome using RNA-sequencing (RNA-seq). Paradoxically, these variants between the samples are usually ignored at the beginning of RNA-seq analyses of many model organisms. This can skew the transcriptome estimates that are used later for downstream analyses, such as expression quantitative trait locus (eQTL) detection. Here, we assessed the impact of reference-based analysis on the transcriptome and eQTLs in a widely-used mouse genetic population: the BXD panel of recombinant inbred lines. We highlight existing reference bias in the transcriptome data analysis and propose practical solutions which combine available genetic variants, genotypes, and genome reference sequence. The use of custom BXD line references improved downstream analysis compared to classical genome reference. These insights would likely benefit genetic studies with a transcriptomic component and demonstrate that genome references might need to be reassessed and improved.

## Introduction

To decipher how genome leads to phenome, measuring gene expression by RNA-sequencing (RNA-seq) is widely used. Fragments of RNA are read and then virtually mapped back onto a reference genome to determine the transcriptomic location of origin. Read mapping is often regarded as trivial but relies on many choices. Indeed, the user decides for example which reference to use and how exact the alignments are required to be. Most of the time, little information is published on how these choices are made. The mapping needs to account for amplification and sequencing errors, and for redundancy within the genome. It is important that the reference precisely represents the samples to guide the mapping. However, generating a reference assembly is complex and expensive and it is common practice to map all samples of a model organism to a single assembly provided by the genome reference consortium (GRC) (Church et al., 2011, 2015). The expression of non-reference alleles may be altered compared to that of reference alleles. This reference bias on the transcriptome can then spread to downstream analyses such as expression quantitative trait loci (eQTL) detection, where gene expression is associated to genomic variants. The genomic variations between individuals at the core of genetic studies are thus paradoxically often ignored at the start of the analysis and may alter interpretations and conclusions.

The genetic characteristics of humans have been widely studied and reference bias is known to alter DNA-seq, RNA-seq (Liu et al., 2018), and chromatin immunoprecipitation (ChIP)-seq analyses (Groza et al., 2020; Rivas-Astroza et al., 2011). The ideal solution would be to use a sample-specific genome assembly. Since this is currently too costly, many methods to reduce reference bias were proposed (Ballouz et al., 2019; Chen et al., 2020; Liu et al., 2018). One strategy notably used by the Genotype-Tissue Expression (GTEx) consortium is to tailor the analysis to each individual as initially implemented in the *WASP* suite of tools (Geijn et al., 2015). The WASP-correction proposes to map reads to the GRC assembly and identify mapped reads that overlap SNVs, then re-map these reads after replacing the reference alleles by variants alleles in the assembly and discard reads that change mapping loci. Although this strategy removes reference bias it also discards reads that are potentially informative of a genetic effect. Nevertheless, the idea of modifying the GRC reference assembly with variants specific to the individual or sample is used by many tools, with the difference that all the reads are mapped to the customized references only. For example, the AlleleSeq pipeline was developed for human trios where the variants for the two parents are known (Rozowsky et al., 2011). One of its tools, *vcf2diploid*, constructs two haplotype-specific references from one reference assembly and a list of genomic variants which can include single nucleotides variants (SNVs), indels, and structural variants (SVs). The authors propose to map the offspring sample separately onto its two parental references, and to retain for each read the alignment with the highest alignment score. In case of equality the alignment is randomly taken from that of either parent to avoid systematic bias. *RefEditor* offers a similar approach, but adds a genotype imputation option (Yuan et al., 2015). Many tools aim at making the best use of large-scale variants and genotypes databases by genotype imputation to have for each individual a more complete set of alleles. However, these genotype imputation strategies cannot be applied to mouse or other animal models because of a lack of genetic characterization at the individual level.

Mouse genetic research mostly uses inbred lines, in which individuals are presumed isogenic. Therefore, it seems logical to aim for reference customization for mouse strains rather than for individuals. The GRC mouse assembly is mainly based on the inbred strain C57BL/6J (B6) (Mouse Genome Sequencing Consortium, 2002) and short genomic variants for many other inbred strains are available at dbSNP (Sherry et al., 2001). To compare retinal transcriptome in two inbred strains (DBA/2J (D2) and B6), Wang et al. modified the GRCm38 reference genome with D2-specific variants from dbSNP to map the D2 samples (Wang et al., 2019). This improved slightly the mappability by reducing the fraction of unmapped reads. *Seqnature* software aims at producing individualized diploid references for RNA-seq analysis and was used on simulated and real world data of Diversity Outbred (DO) mice, in which each mouse is a unique combination between 8 founder strains (Munger et al., 2014). It shows improvement of the number of reads mapped, of the accuracy of transcript expression estimates, and of the number of eQTLs detected. Unfortunately, this type of study is very rare and the R package (DOQTL) used for the QTL analyses is specific to this mouse population, which renders comparisons with studies on different populations challenging. The Mouse Genome Project tries a more global approach to characterize the genetic variation among mouse strains. Many genetic variants were discovered and strain-specific genome assemblies for sixteen mouse strains were released (Lilue et al., 2018). However, it remains unclear how to use these resources for mice that are intercrossed.

The BXD panel of recombinant inbred lines is a well-studied and genetically simple population derived from the B6 and D2 strains (Peirce et al., 2004). Each BXD line has genetic markers (genotypes) available on the GeneNetwork website (http://genenetwork.org). Although this panel has been used in hundreds of studies, nobody to our knowledge has performed neither BXD-specific read mapping, nor BXD genome assembly. Here, we explored different strategies using publicly available resources to accurately represent the genetic diversity of the samples. We assessed the influence of the reference used for read mapping in this panel and how it impacts read mappability, gene expression, and eQTLs. We also measured how various parameter settings would influence the number of eQTLs found. We evaluated the use of the two parental genome assemblies and found this strategy not adequate. We implemented an alternative strategy which enhanced the GRC assembly with known variants. This improved the quality of BXD transcriptomics analyses. Our approach reduces reference bias in the BXD transcriptomics, and raises awareness about pitfalls of RNA-seq analyses.

## Methods

### Samples and RNA-sequencing

Male mice from 33 BXD lines, C57BL/6J (B6), DBA/2J (D2), and F1 (Figure 1A) were either sleep-deprived (SD) for 6 hours starting at zeitgeber time (ZT) 0 (start of lights on) or not (i.e. non-sleep deprived or NSD) (Diessler et al., 2018; Jan et al., 2019). RNA was extracted from cerebral cortex and liver collected at ZT6 in both SD and NSD mice. Prior to sequencing, the RNA was pooled by mouse line and experimental condition (NSD or SD). Single-end reads of 100 bp were obtained using Illumina HiSeq 2500 system. A list of samples, including which BXD lines were used, is available (Table S1).

**Figure 1.**
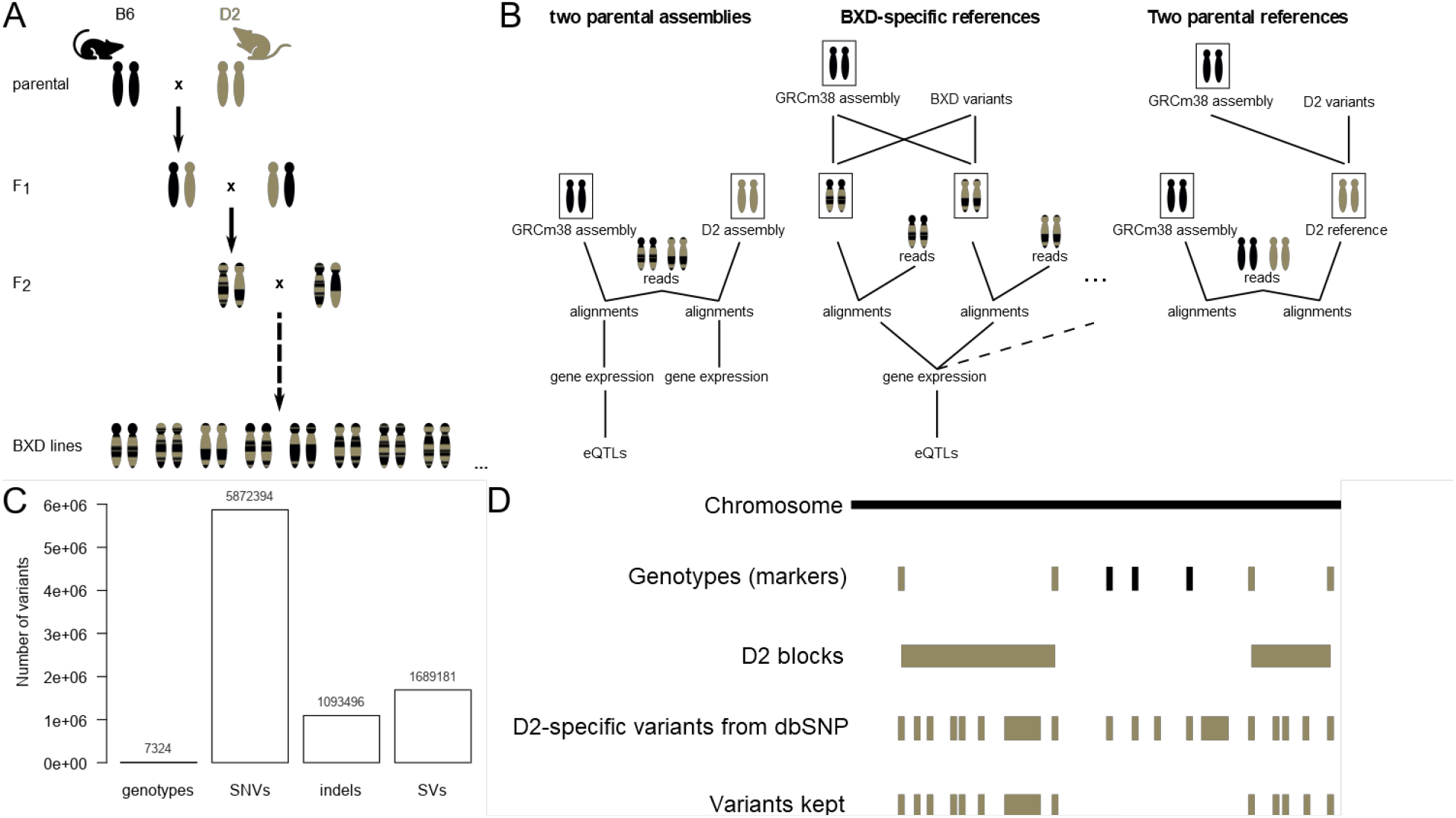
Including genomic variants in inbred mouse transcriptome read mapping overview. A. BXD mouse recombinant inbred panel. Samples come from mice that are: BXD advanced recombinant inbred lines, parental inbred strains which are C57BL/6J (B6) and DBA/2J (D2), and not inbred first generation cross between the parental strains (F1). B. RNA-seq read mapping strategies in this study. C. BXD genotypes available from GeneNetwork and D2-specific genomic variants available from dbSNP. D. Genotypes imputation workflow. B6 regions or alleles are in black, D2 regions or alleles are in brown.

### Genome assemblies and transcriptome annotation

Assembly refers to genomic sequences assembled from DNA reads to form chromosomes, whereas reference simply refers to the sequences the reads are compared to during read mapping. Therefore, an assembly can be used as reference, but different references can be derived from the same assembly. Here, to highlight the generation methods used we reserved the term assembly to external sets of DNA sequences and indicate original/unmodified sequences.

Genome assemblies and transcriptomes annotations are from Ensembl release 94 (ftp://ftp.ensembl.org/pub/release-94). The classical genome sequence is GRCm38 also referred to as mm10 and is based on the B6 strain (Mus_musculus.GRCm38.dna_sm.primary_assembly.fa). The D2 assembly genome sequence was used (Mus_musculus_dba2j.DBA_2J_v1.dna_sm.toplevel.fa) (Lilue et al., 2018). Both genome sequences are not containing alternative haplotypes, and repeats or low complexity regions are soft-masked (sm), which means they are in lowercase letters. The transcriptome annotations are corresponding to these two assemblies (Mus_musculus.GRCm38.94.gtf and Mus_musculus_dba2j.DBA_2J_v1.94.gtf).

### Variants and genotype imputation

The 7324 BXD genotypes available from GeneNetwork are genetic markers selected to be indicative of recombination events between the parental genomes.

(BXD_Geno-19Jan2017_forGN.xlsx, Figure 1C). The D2-specific variants are 5’872’394 SNVs (DBA_2J.mgp.v5.snps.dbSNP142.vcf.gz) and 1’093’496 indels (DBA_2J.mgp.v5.indels.dbSNP142.normed.vcf.gz) from dbSNP (version 142, variants version 5). To have a more complete set of genetic variants specific to the BXD lines, genotype imputation was performed (Figure 1D and Figure S3):

1. The D2 haplotype blocks, defined as sets of at least 2 consecutive genotypes with D2 alleles without B6 or heterozygous alleles in between, were extracted from the GeneNetwork BXD genotypes for each BXD line.
2. dbSNP D2-specific variants were checked for overlap with these D2 blocks using bedtools (version v2.28.0).
3. Variants overlapping with D2 blocks were imputed to be D2 alleles.

During this study, it was noticed that genotypes from GeneNetwork for BXD100 (based on GRCm38 genome assembly also called mm10) had multiple chromosomes without any D2 alleles, which is unexpected considering this was not the case in the previous version of genotypes (based on MGSCv37 genome assembly also called mm9). GeneNetwork has been informed and the error was thought to have occurred during lift-over of the genotypes. We did not try to correct this mistake and kept the erroneous BXD100 genotypes in the current analysis. The effect, if any, on the eQTL analysis should be small, as this is concerning only one BXD lines out of 33, and not all chromosomes are affected.

### Customization of reference

A customized reference genome for each BXD was build based on the GRCm38 assembly and BXD-specific genotypes (from GeneNetwork and imputed). The reference genome sequence GRCm38 was customized for each BXD line with BXD-specific genotypes (from GeneNetwork and imputed) using vcf2diploid software (version 0.2.6) with slight modifications. Prior to compiling the software according to installation instructions, we removed the function that randomizes unphased heterozygous variants (to determine whether there are included in the paternal or maternal genome) and the call to this function, as follows:

**Table 1.** Modifications to vcf2diploid software.

| File name | Code removed |
| --- | --- |
| Variant.java | `public void randomizeHaplotype() { if (_rand.nextDouble() > 0.5) return; int tmp = _paternal; _paternal = _maternal; _maternal = tmp; return; }` |
| VCF2diploid.java | `if (!var.isPhased()) var.randomizeHaplotype();` |

In the modified software, all the unphased heterozygous variants are included in the maternal genome sequence but ignored for the paternal one. We use the paternal sequence so that heterozygous genotypes are ignored. Note that all D2-specific variants from dbSNP and the BXD markers from GeneNetwork are unphased and heterozygous labels may be indicative of low or uncertain quality:

- On GeneNetwork (January 2017), genotypes are defined as “H (heterozygous) if the genotype was uncertain”.
  - 1449 loci have H alleles over the 7320 genotypes (20%) for our 33 BXD lines.
  - Our 33 BXD lines have on average 60 H alleles over 7320 loci and in total 1967 H alleles over 241560 alleles (0.82%).
- For D2-specific variants from dbSNP (version 142), Het means “Genotype call is heterozygous (low quality)”.
  - 481158 SNPs are Het over 5872394 (8%).
  - 80075 indels are Het over 1093496 (7%).

A D2-specific reference genome was build based on the GRCm38 assembly and D2-specific SNPs and indels from dbSNP (Figure 1B “2 parental references”). We refer to this modified version as D2 reference, which is different from the D2 assembly. The coordinates of the transcriptome annotation were adapted to the new coordinates for BXD and D2 references using the chain files generated by vcf2diploid and the liftOver tool (version 8.28) from UCSC (http://genome.ucsc.edu).

### Read mapping and mapping parameters optimization

Read mapping was performed with STAR (version 2.7.0e) (Dobin et al., 2013).

STAR command for default (permissive) alignment:

*STAR --genomeDir genomedirectoryname --outFileNamePrefix output_prefixname*

*--readFilesCommand zcat --readFilesIn reads*.*fastq*.*gz*

STAR command for exact matches (restrictive) alignment:

*STAR --genomeDir genomedirectoryname --outFileNamePrefix output_prefixname*

*--readFilesCommand zcat --readFilesIn reads*.*fastq*.*gz*

*--scoreDelOpen -40 --scoreInsOpen -40 --alignIntronMax 1 --alignEndsType EndToEnd*

*--outFilterMismatchNmax 0*

The transcriptome annotation was not included in the genome index used for exact matches. To count the reads after STAR, HTseq (version 0.6.1p1) was used with samtools (version 1.9) to convert alignments from bam to sam format. Only the alignments with a quality score of 10 or above were kept (default).

*samtools view -h alignment*.*bam* | *htseq-count -s reverse -t exon -m union - reference*.*gtf*

The “-s reverse” parameter is used for the stranded library and is specific to the library preparation and sequencing protocol. Alternatively, for mapping using transcriptome annotation, the HTseq counting implemented in STAR was used using --quantMode GeneCounts.

### Filtering and normalization of gene counts

Lowly expressed genes were filtered by tissue: keeping only genes with counts per million (cpm) above 0.5 (min_cpm) for at least 20 samples (on 66 BXD samples in total). The counts were normalized with edgeR package (version 3.24.3) using the weighted trimmed mean of M-values (TMM) method to take into account the variation in library size and in RNA population (Robinson & Oshlack, 2010) and log transformation (log2).

### Differential mapping analysis of genes

The duplicated gene names were removed, and only gene names common to both GRCm38 and BXD references transcriptome annotation were kept. A differential expression analysis was applied on the BXD samples using voom function from R package limma (version 3.38.3). The two groups of samples compared had only the reference used for read mapping that differed.

### Local eQTL detection and comparison

Note that QTL detection is sometimes referred to as QTL mapping, but we will avoid this terminology to avoid confusion with read mapping. Local eQTLs (often referred to as cis eQTLs) were detected using FastQTL (version 2.184) using 2 Mb bp above and below transcription start site (TSS). 1000 permutations were used to adjust p-values for multiple markers tested and seed 1 was chosen to help reproducibility. Correction for multiple gene testing was performed with R (version 3.4.2) package qvalue (version 2.10.0). The slope (allelic mean difference, representing the direction and strength of allele-specific gene expression) of the linear regression and qvalue of eQTLs from different references were considered similar (unaffected) if they are within less than 5%:

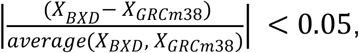

where X refer to slope or qvalue.

### Reference bias

The percentage of skewness of local eQTLs is calculated as:

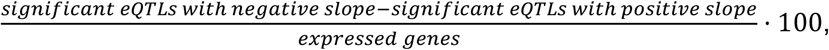

where significant is defined as FDR < 5%.

### Data availability

Raw reads files are accessible on NCBI Gene Expression Omnibus under accession id GSE114845. However, quality information stored in read names is lost upon deposit due to NCBI SRA reformatting, as observed elsewhere (Davis et al., 2021).

Given that data sources are diverse and some do not have a version or identifier, we grouped data on for reproducibility and reusability (10.5281/zenodo.5513980). This repository contains input data: D2-specific variants from dbSNP (vcf), genotypes from GeneNetwork (tab); intermediate and produced data: D2 blocks (bed), imputed variants (vcf), BXD-specific genome sequences (fasta), BXD-specific transcriptome annotation (gtf), gene counts, normalized gene expression, local eQTLs.

### Code availability

The code used for the analyses is on https://github.com/nagobet/BXDmapping.

### Computational requirements

Some computations were performed on the Wally cluster of the University of Lausanne with the Vital-IT software stack (https://www.vital-it.ch) of the Swiss Institute of Bioinformatics for speed (parallelizing multiple mapping runs) and convenience (having a functional installation of FastQTL software). However, none of the steps require unreasonable memory or computational power, and all softwares used in this study are freely available for reproducibility purpose.

## Results

To improve genetic coherency of RNA-seq read mapping, we explored two alternative strategies to exploit available data in the BXD panel. The first strategy uses the two parental strains assemblies (Figure 1B “2 parental assemblies”). The second strategy uses BXD-specific references obtained from the GRC assembly modified with BXD known and imputed variants (Figure 1B “BXD-specific references”). An intermediate between these two strategies was used for comparison: a D2 reference built from the GRC assembly modified with known D2 variants (Figure 1B “2 parental references”). For each strategy, we evaluated the impact on various downstream steps of the analytical pipeline. We looked how the strategies affected mappability of the RNA reads (Figure 2A-C, and Figure 3A&C), gene expression estimates (Figure 2D and Figure 3B), and eQTLs (Figure 4). In addition, we have evaluated how key mapping parameters influence these results (Figure 2B-C, and Figure 5).

**Figure 2.**
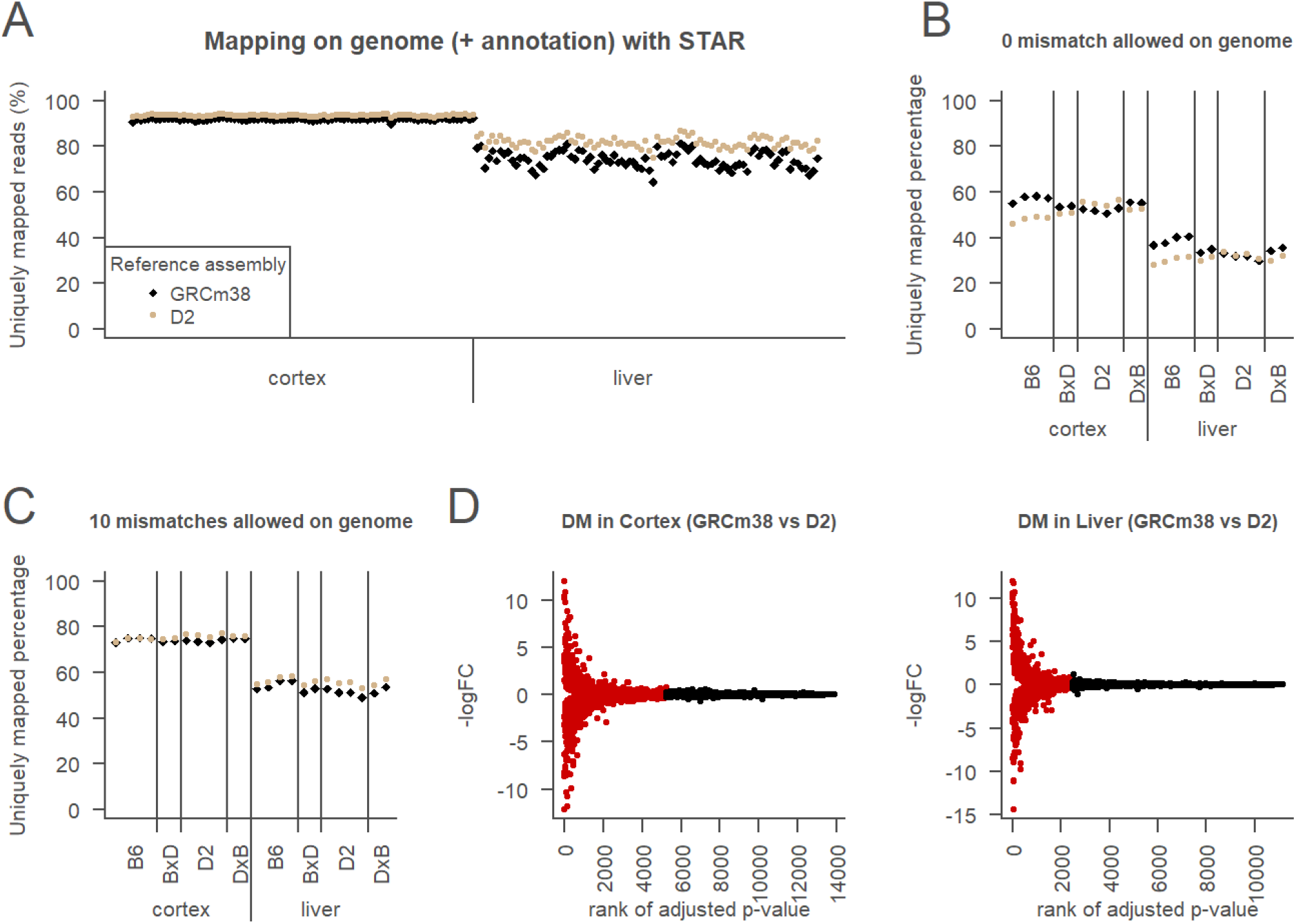
Two parental assemblies strategy. A. Mappability of all samples on 2 parental assemblies (samples are mapped on GRCm38: black symbols and on D2 assembly: brown symbols) using permissive mapping setting (STAR default) in cortex (left) and liver (right). B. Mappability of parental and F1 (BxD and DxB) samples on 2 parental assemblies using restrictive mapping setting allowing 0 mismatches. Same legend than in A. C. Mappability of parental and F1 samples on 2 parental assemblies using restrictive mapping setting but allowing up to 10 mismatches. Same legend than in A. D. Differential mapping (DM) analysis of D2 assembly compared to GRCm38 in the cortex (left) or in the liver (right). Genes are classified as DM genes if FDR adjusted p-value < 0.05 (red) or non DM genes otherwise (black).

**Figure 3.**
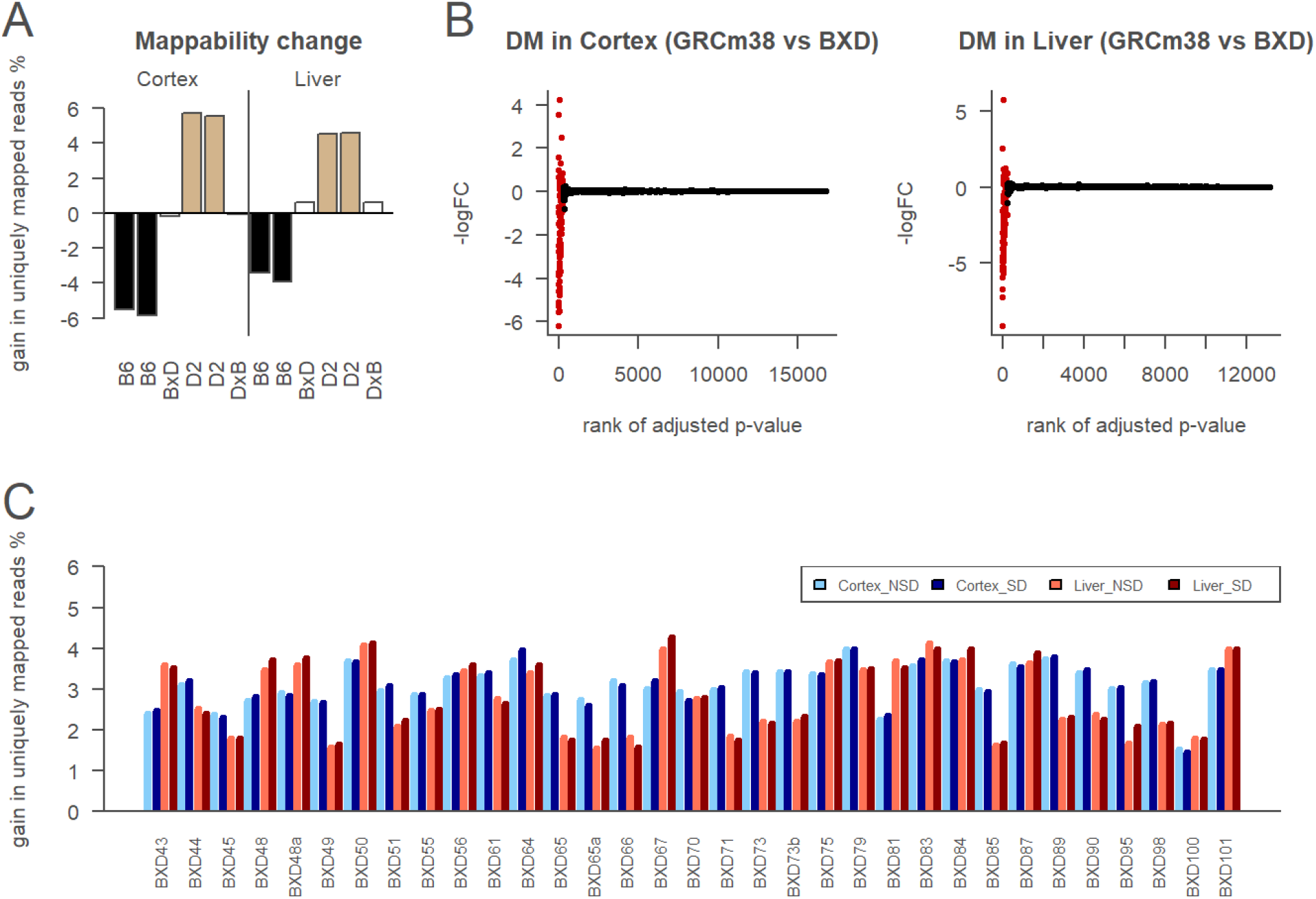
Line-specific references strategy. A. Relative mappability of customized D2-specific reference (GRCm38 modified with D2-specific indels and SNVs from dbSNP) compared to GRCm38 on parental and F1 samples with exact matches. Samples are all NSD. Colors indicate genetic of the samples: B6 (black), D2 (light brown), and F1 (white) between B6 and D2 strains. The F1 samples are BxD if the mother is B6 and the father is D2 (as for the BXD lines), or the reverse for DxB. B. Differential mapping (DM) analysis of customized BXD specific references compared to GRCm38, in the cortex (left) or in the liver (right). Genes are classified as DM genes if FDR adjusted p-value < 0.05 (red) or non DM genes otherwise (black). C. Relative mappability of BXD-specific references (GRCm38 modified for each BXD line with GeneNetwork genotypes and imputed variants) compared to GRCm38 on BXD samples with exact matches.

**Figure 4.**
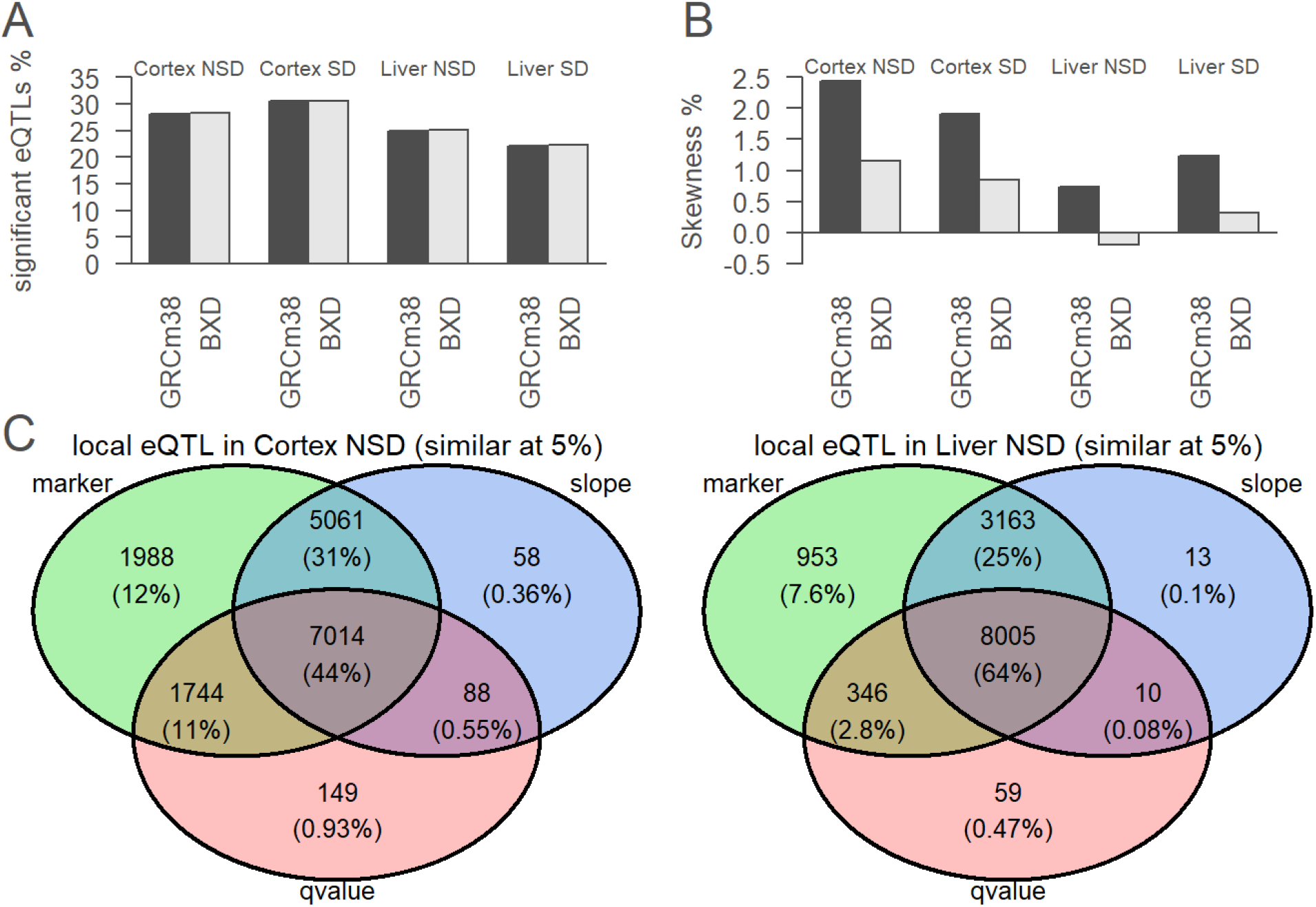
Consequences of mapping reference at local eQTL level. A. Percentage of significant (FDR 5%) local eQTLs with GRCm38 or BXD-specific references. B. Percentage of skewness of significant (FDR 5%) local eQTLs slope with GRCm38 or BXD-specific references. C. Local eQTLs overlapping between GRCm38 and BXD-specific references in cortex NSD (left) or in liver NSD (right). The sign of the slope measures the direction of gene expression (B6 or D2 as more expressed allele).

**Figure 5.**
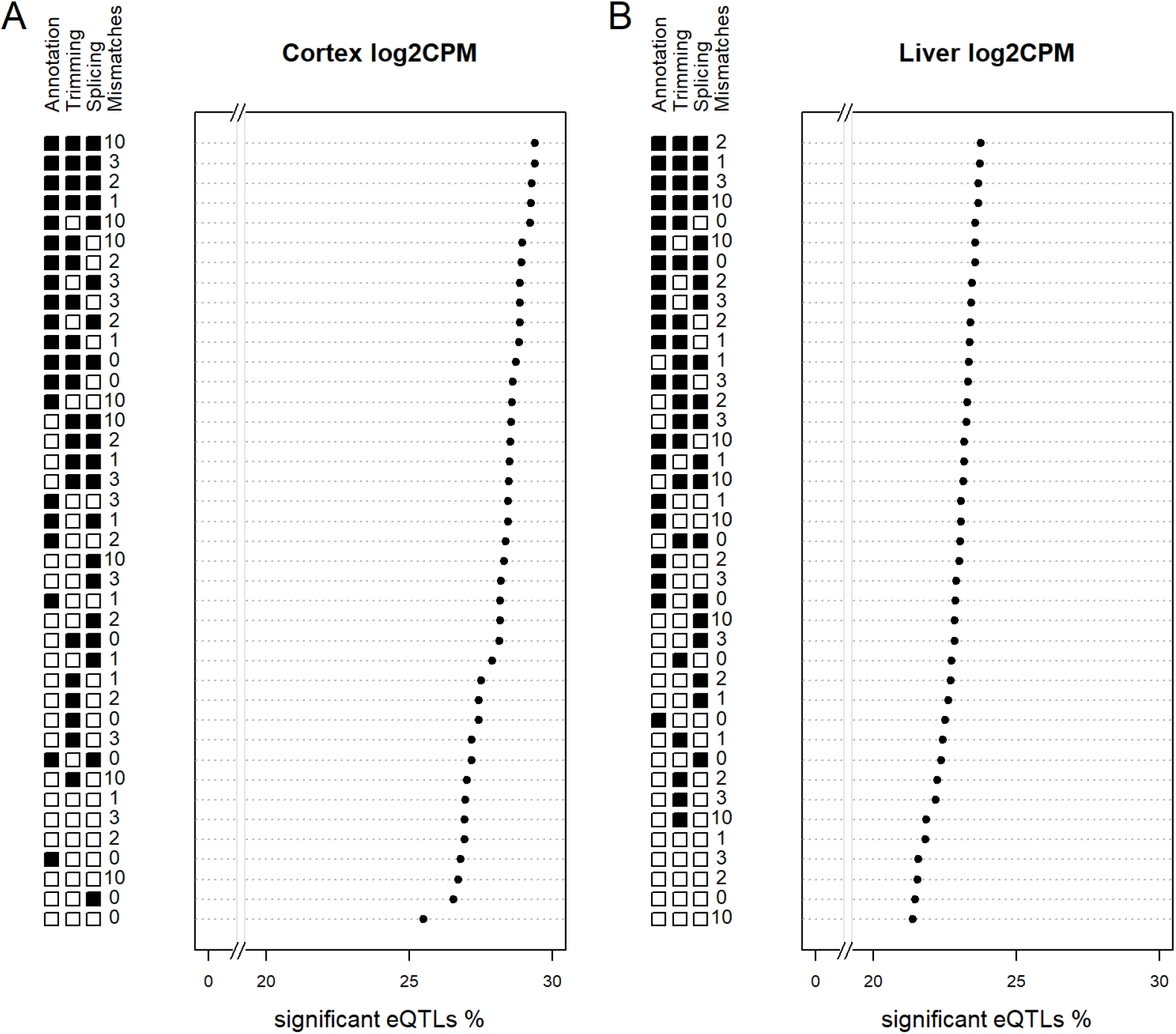
Evaluating mapping parameters. A. The performance on local eQTLs of selected mapping settings on cortex samples (average between NSD and SD conditions) is measured by the percentage of expressed genes than have a significant local eQTLs. B. The performance on local eQTLs of selected mapping settings on liver samples (average between NSD and SD conditions) is measured by the percentage of expressed genes than have a significant local eQTLs.

### Mapping strategy with two parental strains assemblies

To explore the impact of using one reference for all samples despite their genetic differences, we mapped all samples on the classical GRCm38 (B6) genome assembly and on the more recent D2 assembly. We expected that more reads from D2 samples would be uniquely mapped on D2 assembly than on the GRCm38 assembly, and that reads from BXD samples would map approximately equally on both parental assemblies. Surprisingly, we observed that the percentage of uniquely mapped reads, used to estimate mappability, was higher for all samples when mapped to the D2 assembly compared to the GRCm38 assembly (Figure 2A), even for B6 samples. We also noticed that mappability differed between the liver and the cortex both in amplitude and in variance. To further explore the bias for the D2 assembly, we allowed only exact matches. We observed the expected strain-specificity as B6 samples mapped higher on B6 assembly and D2 samples mapped higher on D2 assembly (Figure 2B). Using up to 10 mismatches (the STAR default for 100 bp reads) but no insertions, deletions and trimming, we lost strain-specificity (Figure 2C). However, the more restrictive mapping setting also importantly reduced the number of uniquely mapped reads (Figure 2A-C). This raised the question what choice of parameter settings ensures both high read yield and strain specificity (see part “Mapping parameters evaluation” below).

To determine the impact of mapping reference on gene expression, we performed a differential mapping (DM) analysis. The principle is the same as differential expression analysis, but the mapping references are compared instead of different groups or perturbations. Note that the reads and the values used for the mapping parameters are identical for both references, so differences observed are caused strictly by the reference. More than one third (38%) of genes were affected by the mapping reference in the cortex and about a quarter (22%) in the liver (Figure 2D). However, alignments of the top 4 highly affected genes were visually inspected and revealed variation in the quality of the assemblies and their transcriptome annotation at these precise places. What appeared as differences in gene expression between the two assemblies could be artefacts and not consequences of genetic variants (two examples of artefacts in Figure S2).

### Customizing reference for D2 and BXD lines

To avoid differences of quality and completeness between B6 and D2 assemblies, we modified the B6 reference assembly using SNPs and indels specific to the D2 strain from dbSNP. We mapped parental and F1 samples on these two assemblies with exact matches. The percentage of uniquely mapped reads was increased when the samples were mapped to their corresponding strain reference, compared to the other parental strain reference (Figure 3A). Indeed, D2 samples gained between 4.6 and 5.7% when mapped to the customized D2 reference, whereas B6 samples lost between 3.4 and 5.8%. In contrast, when mapped on the D2 assembly (Figure 2B) D2 samples gained between 0.0% and 3.4% whereas B6 samples lost between 8.5% and 9.8%. The D2 customized reference appears more balanced. In both cases, the difference between the two parental references was smaller for F1 samples, which is expected for a mix between the two parental strains. To apply the same strategy to BXD samples, genotypes were imputed using the large amount of D2-specific variants from dbSNP and the BXD specificity of genotypes from GeneNetwork. All BXD samples gained between 1.4 and 4.3% from having a customized reference by BXD line (Figure 3C). The amplitude of the gain varied among BXD lines and between tissues with cortex samples having globally higher values than liver samples. Note that it was expected for the gain to be lower for BXD samples than for D2 samples, since in BXD lines approximately half of the alleles are D2.

To determine the impact of reference customization on gene expression, we performed a differential mapping analysis. Around 2% of genes are affected by the mapping reference in the cortex and in the liver (Figure 3B).

### Consequences of customization on local eQTL detection

To evaluate the effect of reference customization on estimated biological phenotypes by downstream analysis, we detected local eQTLs using gene expression estimated with the B6 reference or the BXD-specific references. The eQTLs are particularly likely to be influenced since they correlated gene expression to genetic variants. The percentage of significant eQTLs was slightly higher (0.1% difference) when using BXD-specific references than when using B6 assembly (Figure 4A). However, this does not necessarily mean that the eQTLs detected are the same. They can differ by the genetic marker, the direction of gene expression (B6 or D2 as more expressed allele), or change in the q-value (Figure 4C and Figure S4). When considering these 3 variables, mapping reference did not affect 44% of local eQTLs in the cortex and 64% in the liver.

### Reference bias

Next, we wanted to detect a potential reference bias, where B6 alleles get more easily mapped than D2 alleles or the contrary. In DNA-seq studies, this can be achieved by checking the symmetry of the distribution of allelic ratios at heterozygous loci. In RNA-seq, allele-specific expression can also modify allelic ratio. However, we assumed that globally B6 alleles are equally expressed than D2 alleles. Moreover, since our samples are inbred lines, the heterozygous sites are greatly rarer than in other populations (and likely of lower quality), so we compared samples with different homozygous alleles, rather than heterozygous alleles from one sample. The percentage of skewness represents how many local eQTLs deviate from a situation without reference bias (skewness 0%). A positive percentage indicates a B6 bias: more eQTLs with the B6 allele increasing gene expression, whereas a negative percentage indicates a D2 bias: more eQTLs with the D2 allele increasing gene expression. Using the B6 reference shows a reference bias for B6 alleles in all tested tissues and conditions while using BXD-specific references decreased bias, and not always for B6 alleles (Figure 4B).

### Mapping parameters evaluation

The reference used is not the only thing influencing read mapping. To test which values to use for the more critical mapping parameters of STAR we varied: i) the number of mismatches allowed, ii) the possibility to trim end of reads, iii) the possibility to splice reads, and iv) the use of known transcriptome annotation. The ratio of eQTLs to expressed genes was used as an evaluative measure to maximize (Figure 5), as others in eQTL field (Munger et al., 2014; Rubinacci et al., 2021). The idea behind it is that eQTLs can be regarded as allele-specific structured gene expression, and thus more likely to result from actual allele differences than from unstructured general noise. The best settings in both tissues are to use trimming, splicing, and transcriptome annotation. However, the maximal number of mismatches differed: 10 mismatches in the cortex (Figure 5A), but only 2 in the liver (Figure 5B). All top settings use existing transcriptome annotation and thus appears to be the more important parameter.

## Discussion

Genomic variations among individuals are the core of genetic studies. Yet it is common practice in the field to use one assembly as reference for all genetically different samples. Here, we improved genetic specificity of read mapping of BXD samples using publicly available data. Our custom BXD-specific references detected proportionally more eQTLs and alleviated reference bias. Below we will discuss the complexity of assessing the analytical design of RNA-seq and the various strategies to integrate genomic variations in transcriptomic analyses.

Although the analysis of RNA-seq data is often regarded as well established, it remains a complex procedure. For mapping of real samples, the true location of reads is unknown. The fraction of uniquely mapped reads is used as a mapping statistic because an RNA molecule can only come from one locus. However, this does not guaranty the correctness of mapping of uniquely mapping reads because, due to e.g. redundancy in the genome, some reads are expected to be correctly classified as multi-mappers. Moreover, uniqueness can have slightly different meanings depending on the mappers and parameters used, as reads are not necessarily exactly and fully aligned because of mismatches, indels, and sequencing errors especially at the ends of reads (trimming).

Most studies are using standard analytical pipelines with default setting, or with slight modifications such as the number of mismatches allowed. However, even though it has been shown to importantly impact genetic specificity (Yuan et al., 2015), the choice of the number of mismatches allowed is rarely given. We showed that the mapping setting has an effect and that surprisingly, the optimal number of mismatches allowed seems to differ between the two tissues tested. We were unable to identify supportive literature for this phenomenon. We think that a more thorough analysis of such tissue effects (e.g. in GTEx) is warranted. An interesting benchmark was completed on human RNA-seq data in a differential gene expression (DE) pipeline (Sha et al., 2015). They compared filtering methods, along with transcriptome reference sets, normalization and DE detection, alignment and counting software. They concluded that the optimal filtering threshold depended on the pipeline parameters. Particularly, the mapping software had the least impact whereas the transcriptome annotation had the higher impact.

Also, our study highlighted the importance of transcriptome annotation even if our evaluating measure differed. Our study focused on the impact of genetic differences in the reference and assessed different combinations of mapping options. In our case, the fraction of eQTLs was used as evaluation measure rather than DE, because we compare a genetic population and not two fixed groups. Indeed, each BXD sample has both B6 and D2 alleles, so the samples compared for each gene are not split the same way, depending on alleles at this locus. A reference bias is likely to influence eQTLs because a variant in a certain gene can modify the mapping of the reads precisely for the samples that have the alternative allele in this region.

An assembly is a global solution because it uses all genomic variants specific to a strain regardless of their size. However, we showed that the gap in quality and completeness between the two parental assemblies is masking the genetic specificity. In addition, the transcriptome was more studied in B6. Even if transcriptome annotation is likely to be similar in other strains, the different coordinate systems between the assemblies complicate the transfer. Moreover, the actual D2 transcriptome annotation corresponding to the D2 assembly includes manual curation steps that make it hard to update to new releases of the genome assembly or variants. Notably, no study was published using this D2 assembly, except the group that released it. In contrast, our customization of one assembly offers the advantage that the coordinate changes are formalized which allows automatization of transcriptome annotation changes with the same tool used for upgrading versions of an assembly (liftOver). For mapping reads of D2 samples and those of other strains than B6, we currently recommend the use of GRCm38 assembly modified with strain-specific indels and SNVs from dbSNP.

Our custom references combine the specificity of BXD genotypes with the large amount of D2-specific short variants from dbSNP. Importantly, structural variants (SVs) are not included, although many were detected between B6 and D2 strains (Quinlan et al., 2010). SVs can have important phenotypic impacts (Mahmoud et al., 2019), potentially more than SNVs (Keane et al., 2011; Scott et al., 2021). However, SVs calls require further efforts in the reporting to ensure the confidence and the format for integration into current workflows. This is due largely to the nature of SVs: their length and large variety implies that the possible number of SVs is greatly superior to that of SNVs, making them less easy to validate and report. Without technologies like long read, optical genome mapping, those SVs will be very likely inaccessible for mouse models unless an international consortium tackles this issue.

Another limitation is that all murine assemblies are haploid whereas mice are diploid. The diploidy is ignored at the mapping step under the assumption that the genotypes of inbred strains are mostly homozygous. However, the homozygosity and stability of inbred mouse strains is based on a theoretical model without occurrence of new mutations (Casellas, 2011), but experimental estimations support a non-neglectable rate of mutations (Casellas & Medrano, 2008), with between 10 and 30 germline mutations occurring per generation (Reardon, 2017). Unfortunately, the assumption of stability of inbred lines is so strongly anchored in the field that its verification is compromised, because of not searching for heterozygous sites or dismiss them. Indeed, the term heterozygous is sometimes used to call variants uncertain or low quality, and they are always unphased. When mouse assemblies were built, regions with high density of heterozygous sites were used to detect (haploid) assembly errors, ignoring the potential coherence of diploid or polyploid references (Lilue et al., 2018). More systematic detection and characterization of heterozygous regions will likely improve the accuracy of transcriptomics studies, particularly for loci with allele-specific expression. However, the read mapping of different possible alleles, which could also be used for not inbred crosses between 2 strains would require a reconciliation step, as implemented for example for human with paternal and maternal allele (Rozowsky et al., 2011). Indeed, every read can come from one allele or another but not from both at the same time. We made efforts to improve the D2 parts of reference to map BXD samples, however the B6 strain itself is also susceptible to mutations and genetic shift, as confirmed by the occurrence of substrains. It is likely that B6 samples used to build the GRC assembly and the B6 samples used currently present some genetic drift. Therefore, a complete characterization of genetic variants of the BXD by DNA-sequencing could improve the customization of both D2 and B6 parts of BXD, and therefore enhance resolution of downstream analyses.

## Conclusion

In current genetic studies using the BXD population, genomic variations are paradoxically ignored at the read mapping step, which as we show here causes a reference bias. The genomic variations need to be explicitly integrated in the reference instead of treated as sequencing errors. Our results show the need for a critical evaluation of the RNA-seq pipeline and the development of more complete genomic variants databases to best approximate the genetics of the samples. Most genetic studies with a transcriptomic component in mice and other model organisms can suffer from reference bias, which could be attenuated by assessing and sequencing those strains. The mouse community could follow the drosophila community (http://dgrp2.gnets.ncsu.edu) and sequence genetic reference populations. Our study can serve as a wake-up call for improving the characterization of genomic variations, and as a concrete guide for analyses in BXD and other genetic populations. As RNA-seq analyses are often a starting point to identify one or a few genes that then are studied in more detail in follow-up experiments, it is worth the extra effort to avoid potential bias by not blindly following traditional pipelines.

## Acknowledgements

We thank Mark Ibberson and Brian Stevenson for the helpful discussions and guidance. P.F. and M.J. were funded by the University of Lausanne (Etat de Vaud). N.G. was funded by the Swiss National Science Foundation grant to P.F. (31003A_173182 and 310030B_192805). I.X. is funded by the CHUV/UNIL (Etat de Vaud).

## Authors contributions

M.J. had the initial idea of the project. N.G., M.J., P.F., and I.X. conceived the experiments. N.G. implemented the experiments, generated figures, and wrote manuscript with feedback from all authors. M.J., P.F., and I.X. supervised the project.

## Supplementary Information

TableS1_Samples.csv

**Table S1** | **Mouse replicates**.

**Figure S1.**
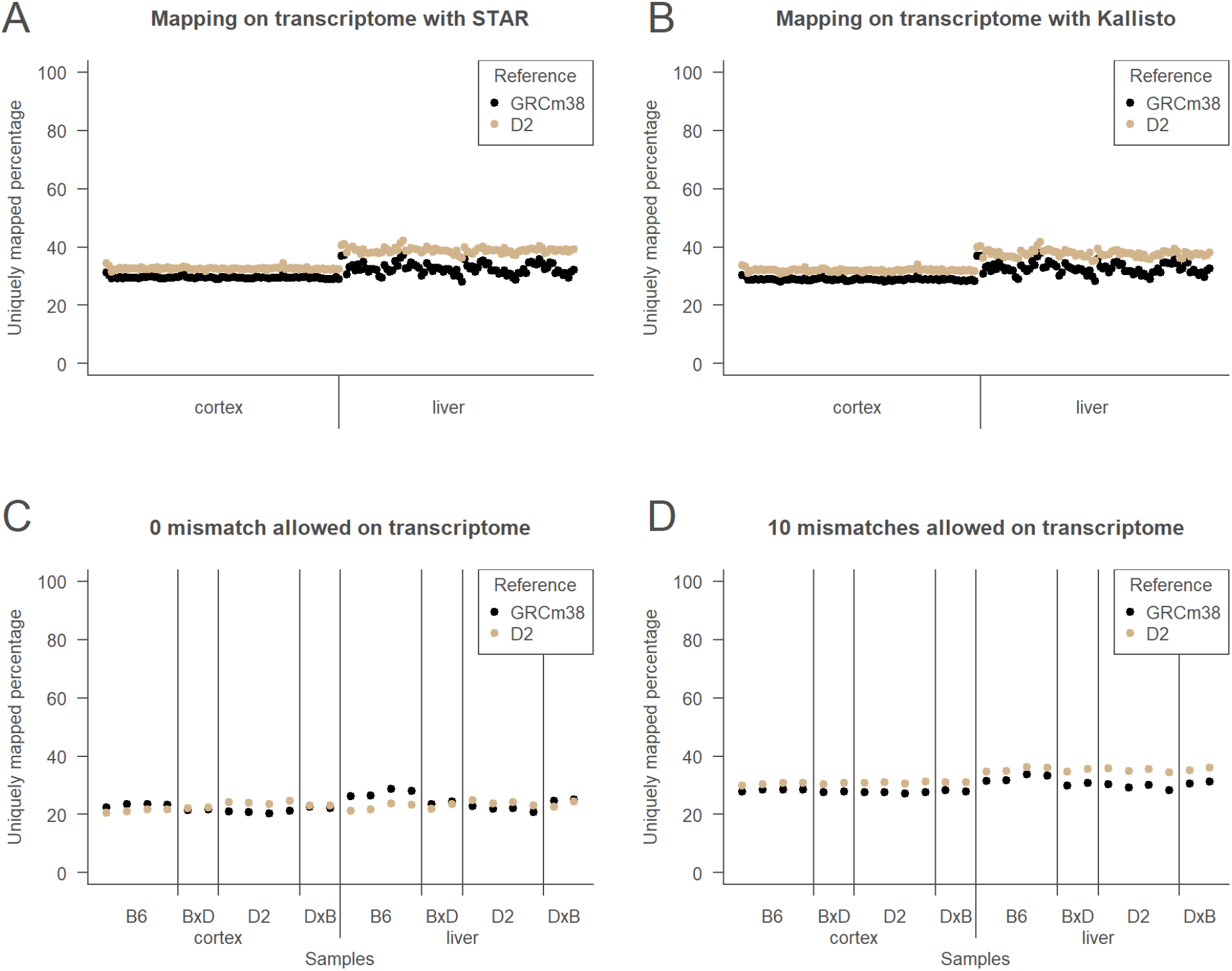
Consequences of mapping reference transcriptome at read mapping level. A. Mappability of all samples on 2 parental assembly transcriptomes using STAR permissive mapping setting. B. Pseudo-mappability of all samples on 2 parental assembly transcriptomes using Kallisto. C. Mappability of parental and F1 samples on 2 parental assembly transcriptomes using STAR restrictive mapping setting. D. Mappability of parental and F1 samples on 2 parental assembly transcriptomes using STAR restrictive mapping setting, but up to 10 mismatches.

**Figure S2.**
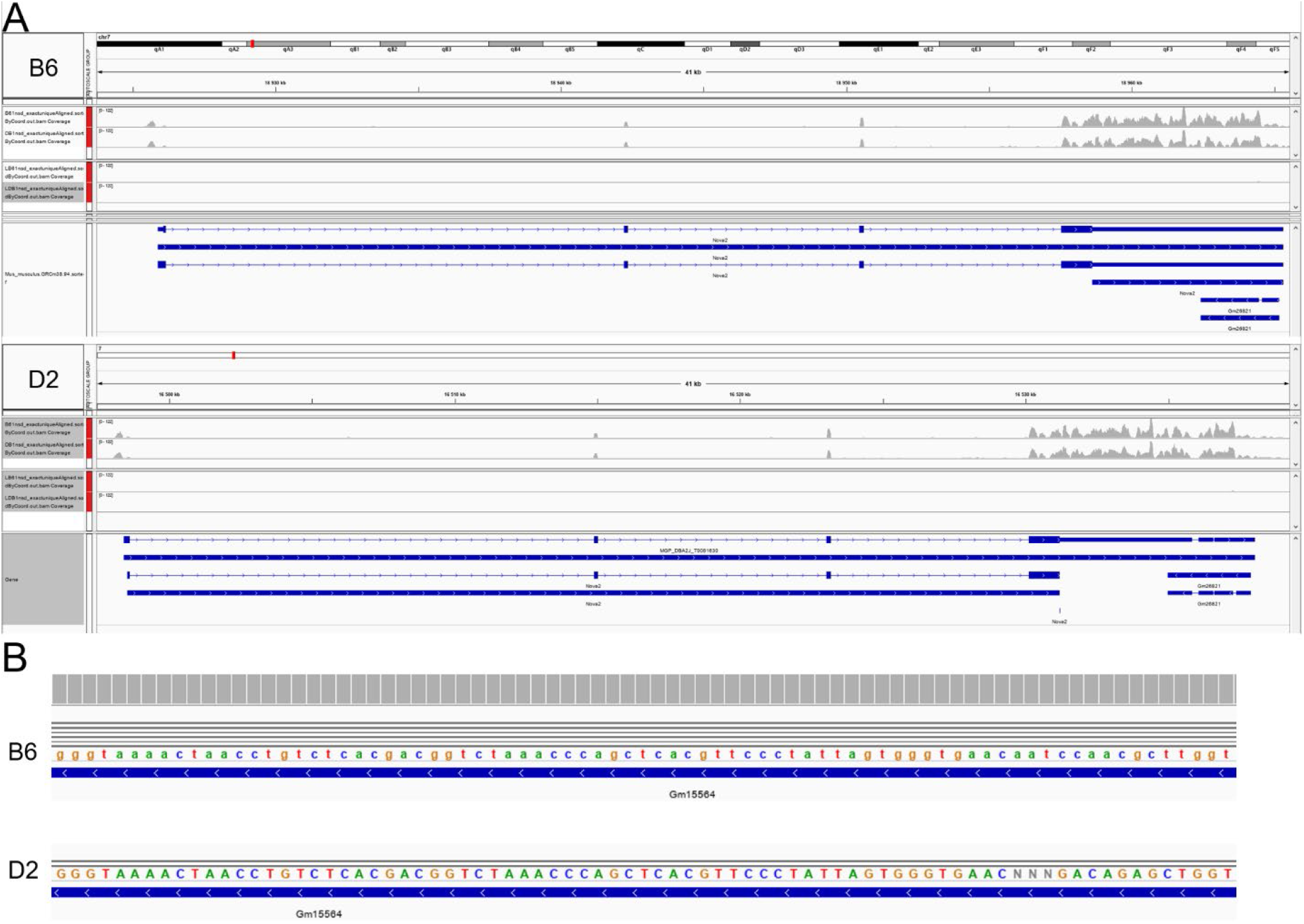
Examples of artefacts of assemblies sequence and annotation. *A. Nova2* genomic region in Integrative Genomics Viewer (IGV), as an transcriptome annotation artefact. The coverage is very similar between B6 and D2 assemblies, but the annotation differs, which causes the reads to be counted differently. *B. Gm15564* genomic sequence in IGV, as an artefact due to difference in completeness of genome assembly. Many reads map to this region on B6 assembly, but not on D2. It appears that in this region of the D2 assembly there are three unknown nucleotides (with label “N”), which supports the interpretation that it is probably due a difference in assembly quality, and not to a genomic variant.

**Figure S3.**
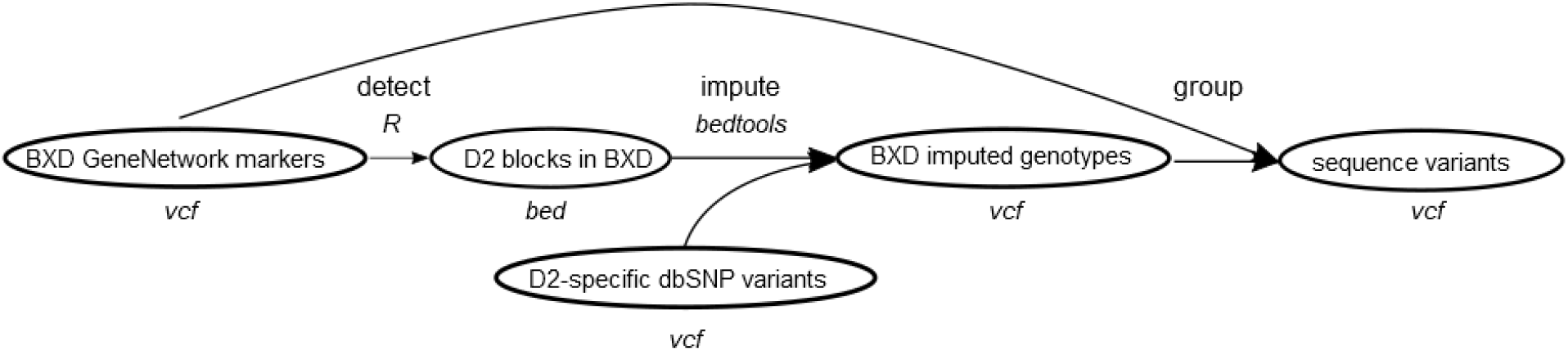
Genotype imputation workflow.

**Figure S4.**
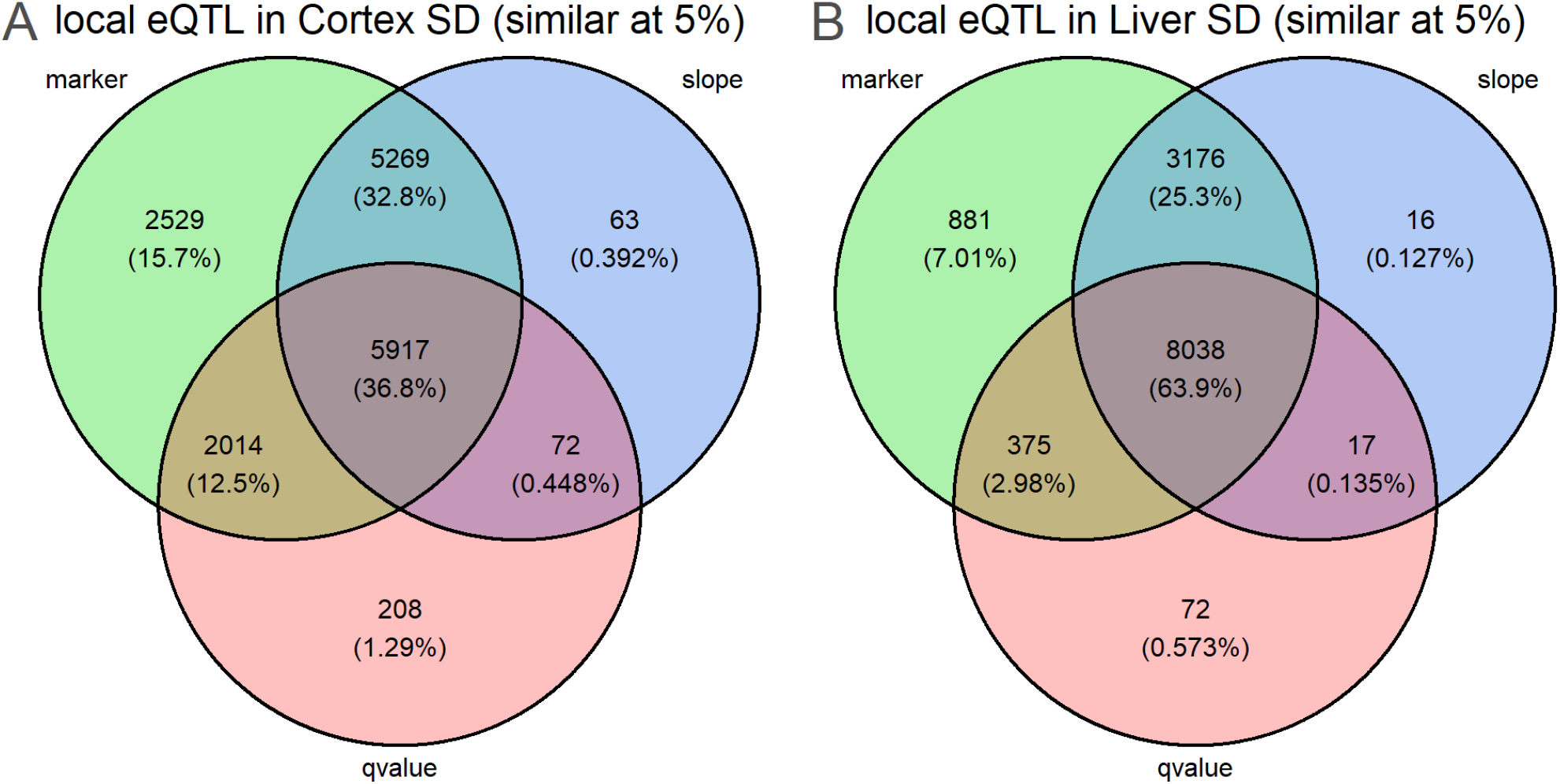
Consequences of mapping reference at local eQTL level in SD condition. A. Local eQTLs overlapping in cortex SD between GRCm38 and BXD-specific references. B. Local eQTLs overlapping in liver SD between GRCm38 and BXD-specific references.

